# Quantification and Localisation of New Brain Lipid Synthesis Using Deuterium Oxide and High Resolution Mass Spectrometry

**DOI:** 10.64898/2025.12.14.694259

**Authors:** Catherine Zhang, Jesse A. Michael, Jonathan D. Teo, Huitong Song, Mika T. Westerhausen, Shadrack M. Mutuku, Shane R. Ellis, Anthony S. Don

## Abstract

Myelin is the lipid-rich membrane that surrounds neuronal axons and is essential for neurological function in vertebrates. The development of therapeutics that stimulate myelin repair to treat demyelinating disorders such as multiple sclerosis is hampered by the inability to distinguish newly-synthesised from pre-existing myelin. This study aimed to develop a method to quantify and localise new myelin lipid synthesis *in vivo*. Deuterium oxide was administered for two weeks in the drinking water of mice fed normal chow, chow containing the demyelinating toxin cuprizone, or during spontaneous remyelination following cuprizone withdrawal. Liquid chromatography-tandem mass spectrometry and mass spectrometry imaging were used to quantify and localise the newly synthesised, deuterated lipids. While most glycerophospholipids were constitutively deuterated, deuteration of myelin-enriched sulfatides, hexosylceramides, and phosphatidylethanolamine plasmalogens was only apparent during remyelination. Deuterated hexosylceramide and phosphatidylethanolamine plasmalogen species were localised primarily to the corpus callosum, the white matter tract that is most heavily affected by cuprizone. Most deuterium atoms were found in the fatty acyl chains, indicative of *de novo* lipid synthesis. These methods provide the means to quantify and spatially profile dynamic lipid synthesis across diverse biological contexts, including understanding myelin homeostasis and preclinical evaluation of remyelinating therapeutics.

## Introduction

Myelin is a lipid-rich membrane produced by specialised cells called oligodendrocytes in the central nervous system, and Schwann cells in the peripheral nervous system. It is wrapped in a spiral fashion around neuronal axons, and is required for the rapid conduction of electrochemical signals along axons.^[1]^ Demyelinating diseases such as multiple sclerosis (MS) have debilitating consequences including vision loss, muscle weakness, and cognitive dysfunction.^[2]^ Current, clinically approved MS therapeutics are effective at preventing immune-mediated demyelination, however remyelination is limited, preventing functional recovery. A stronger understanding of factors that modulate remyelination and the development of pro-myelinating therapeutics will improve outcomes for people with demyelinating diseases.^[3]^

Assessing remyelination *in vivo* is challenged by the inability to identify and quantify newly-formed myelin. Measurements of myelin density and thickness using histological staining and electron microscopy quantify total myelin content at a given point in time and cannot differentiate newly-formed from pre-existing myelin, yet are widely used to assess remyelination.^[4,5]^ Magnetic resonance imaging can be used to visualise myelin content in living organisms, but similarly cannot distinguish new from pre-existing myelin. Transgenic mice expressing a fluorescent protein in newly formed oligodendrocytes provide a surrogate measure of new myelin,^[6]^ and have been used to follow remyelination *in vivo*.^[4,7]^ Although useful, this approach is semi-quantitative, does not quantify actual myelin constituents, and assesses myelin synthesis only by newly-formed (as distinct from existing) oligodendrocytes. Additionally, this approach requires complex transgenic models.

Myelin is 70-80% lipid (dry weight), of which 20-25% is hexosylceramide (HexCer) and 5% is hexosylceramide-sulfate (sulfatide, SHexCer).^[8,9]^ In the central nervous system, over 99% of HexCer and SHexCer are galactosyleramides.^[10,11]^ These lipids are synthesised exclusively by oligodendrocytes and hence unique to myelin.^[12-14]^ Myelin is also enriched in cholesterol and certain phosphatidylethanolamine plasmalogens (PE-P).^[9,15]^ Developments in liquid chromatography-tandem mass spectrometry (LC-MS/MS) over the last two decades have enabled high throughput analysis and quantification of hundreds of lipids in any given sample. Added to this, mass spectrometry imaging (MSI) now permits the localisation of lipids with low to sub-micron resolution.^[16-18]^ However, conventional lipidomic analysis and MSI do not discriminate between newly-synthesised and pre-existing lipids.

Deuterium oxide (^2^H_2_O) administration provides a means to follow new lipid synthesis in a defined time frame through metabolic incorporation of deuterium (^2^H) during endogenous lipid synthesis.^[19] 2^H_2_O can be administered by mixing with drinking water, rapidly equilibrates with body fluid, and is safe at levels up to 30% in the drinking water of mice.^[20-22]^ During *de novo* lipogenesis, ^2^H is incorporated into fatty acids chains through ^2^H-labeled acetyl-CoA and malonyl-CoA and into the glycerol backbones of triglycerides during glycolysis and glyceroneogeneis.^[23,24] 2^H incorporation into fatty acids can also occur through enzyme-catalysed hydrogen-deuterium exchange from ^2^H-labeled NADPH.^[24]^ Early studies used ^2^H_2_O in combination with gas chromatography-mass spectrometry to investigate the turnover of specific fatty acids, cholesterol, and triglycerides in animals and humans.^[25]^ Using pre-fractionation of lipid classes coupled to GC-MS analysis of ^2^H-labelled palmitate, Ando *et al*. demonstrated that myelin cholesterol and galactosylceramide are turned over much slower than glycerophospholipids, and that their turnover is affected by age.^[26]^ More recently, Goh *et al*. applied ^2^H_2_O in combination with LC-MS/MS lipidomic analysis to determine the fractional synthesis of approximately 100 lipids from 13 functional classes in cultured cells.^[27]^ This approach was later adapted by Baumann *et al*., who administered ^2^H_2_O to pregnant mouse dams and compared the levels of deuteration for 26 lipids in developing versus adult neural tissue using LC-MS.^[28]^

In this study, we demonstrate that ^2^H_2_O administration in the drinking water can be used in combination with LC-MS/MS and MSI to quantify and visualise new myelin synthesis in a mouse model of toxin-induced demyelination. Cuprizone is a copper-chelating toxin that induces oligodendrocyte death and demyelination in mice.^[29]^ Spontaneous remyelination occurs within two weeks after cuprizone withdrawal.^[30]^ LC-MS/MS analysis of corpus callosum tissue revealed consistently high levels of deuteration in most glycerophospholipids, indicative of continuous turnover. In contrast, significant deuteration of myelin-enriched HexCer, SHexCer, and PE-P species was only apparent during remyelination following cuprizone withdrawal. MSI revealed localisation of these deuterated lipids to major white matter tracts, particularly the corpus callosum. This approach can be broadly applied to monitor and quantify specific, localised lipid metabolism events *in vivo*.

## Results and Discussion

### Quantification of Deuterated Lipids in the Corpus Callosum

To quantify new lipid synthesis during demyelination and remyelination, mice were administered 25% ^2^H_2_O in their drinking water for the final two weeks of a five-week cuprizone diet (Cpz), or for two weeks after removal of cuprizone from the diet (Rem) (Figure 1a). A control group received only normal mouse chow and 25% ^2^H_2_O for two weeks (Ctrl) to measure baseline lipid synthesis. As expected, cuprizone feeding caused significant myelin loss in the corpus callosum, and spontaneous remyelination was apparent two weeks after cuprizone withdrawal (Figure 1b). Remyelination was more notable in the posterior compared to the anterior corpus callosum.

**Figure 1.**
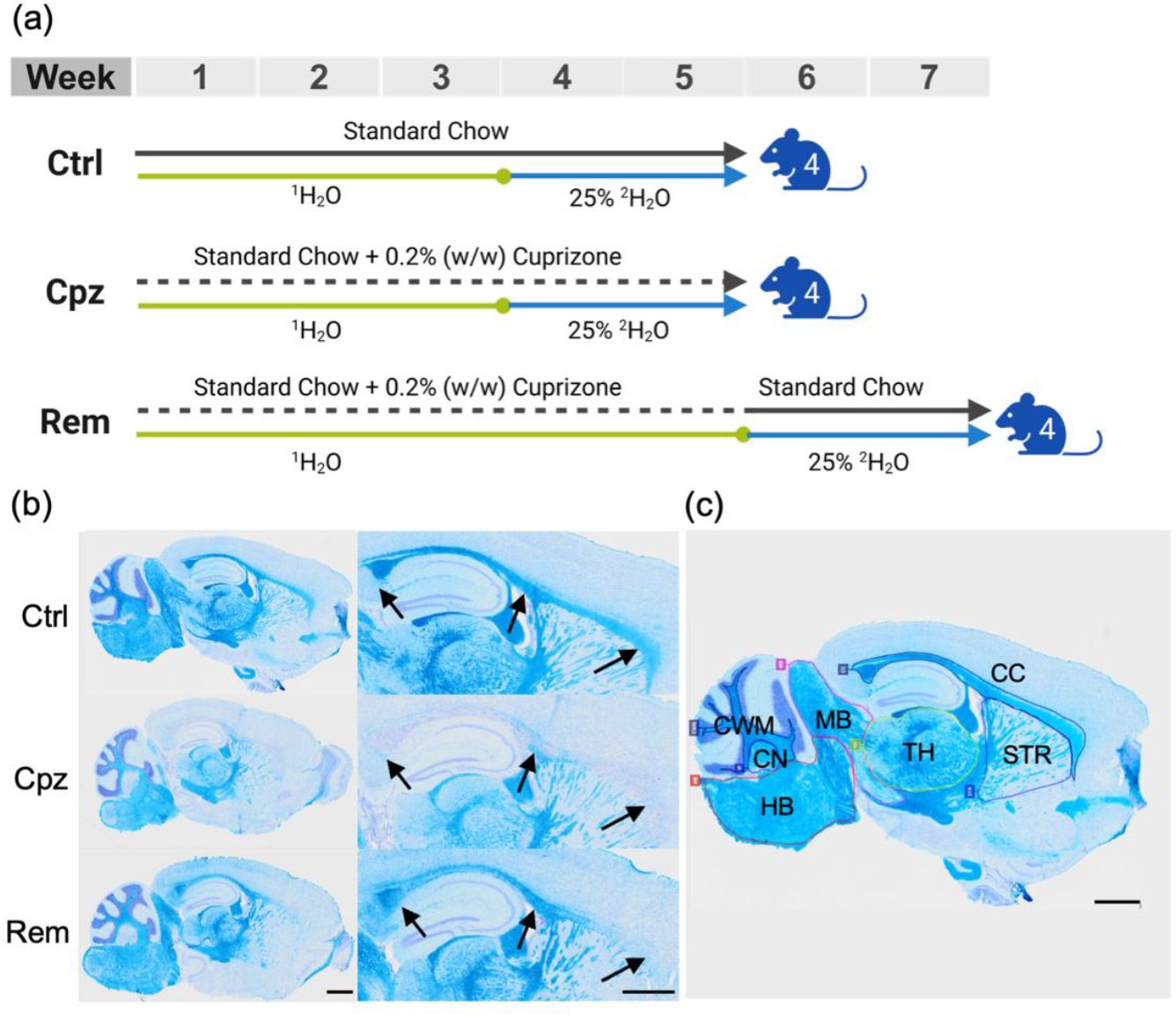
Experimental design. a) Schematic of the experimental design. Mice were provided 25% ^2^H_2_O in their drinking water (^1^H_2_O) for the final two weeks of a five-week cuprizone diet (Cpz, n = 4), or for two weeks after removal of cuprizone from the diet (Rem, n = 4). The control group received only standard chow and 25% ^2^H_2_O for the final two weeks (Ctrl, n = 4). An additional group of mice (not shown) was provided ^1^H_2_O throughout the entire duration of each diet (n = 1 each for Ctrl and Cpz, n = 2 for Rem). b) Representative images of luxol fast blue staining of myelin in sagittal brain sections from each experimental group. Cell nuclei are counterstained with cresyl violet in purple. Scale bar: 1000 μm. Cuprizone induced prominent demyelination in the corpus callosum (indicated by arrowheads), and remyelination was apparent two weeks after cuprizone withdrawal. c) Annotation of brain regions. From left to right: cerebellar white matter (CWM), cerebellar nuclei (CN), hindbrain (HB), midbrain (MB), thalamus (TH), Corpus callosum (CC), and striatum (STR).

The workflow for quantification of deuterated lipids is shown in Figure 2a. Lipid identifications were assigned to 171 LC-MS/MS peaks, comprising the most abundant lipid species in each of 16 lipid classes, based on MS/MS spectra in lipid extracts of corpus callosum tissue from mice administered drinking water without ^2^H_2_O (Table S1, Supporting Information). LC-MS/MS peaks corresponding to deuterated isotopologues (M+1 – M+20) were then identified in corpus callosum lipid extracts from mice administered ^2^H_2_O, using a combination of accurate precursor *m/z (*10 ppm mass tolerance) and elution time within 0.1 min of the non-deuterated monoisotope (M+0) (Figure 2b). To eliminate the effects of naturally occurring heavy isotopes on the quantification of deuterated isotopologues, the isotope correction algorithm IsoCor was applied to correct the peak areas based on the natural abundance of heavy isotopes.^[31]^ Lipids were considered to show quantifiable deuteration if (i) the abundance of deuterated isotopologues relative to the M+0 peak (after isotope correction) was below 5% in the ^1^H_2_O-only control samples; and (ii) four consecutive isotopologue peaks were obtained in a minimum of three out of four samples in any of the three groups receiving ^2^H_2_O (Figure 2c, d). The complete lipidomic dataset containing the quantified amounts of lipid isotopologue is presented in Table S2 (Supporting Information).

**Figure 2.**
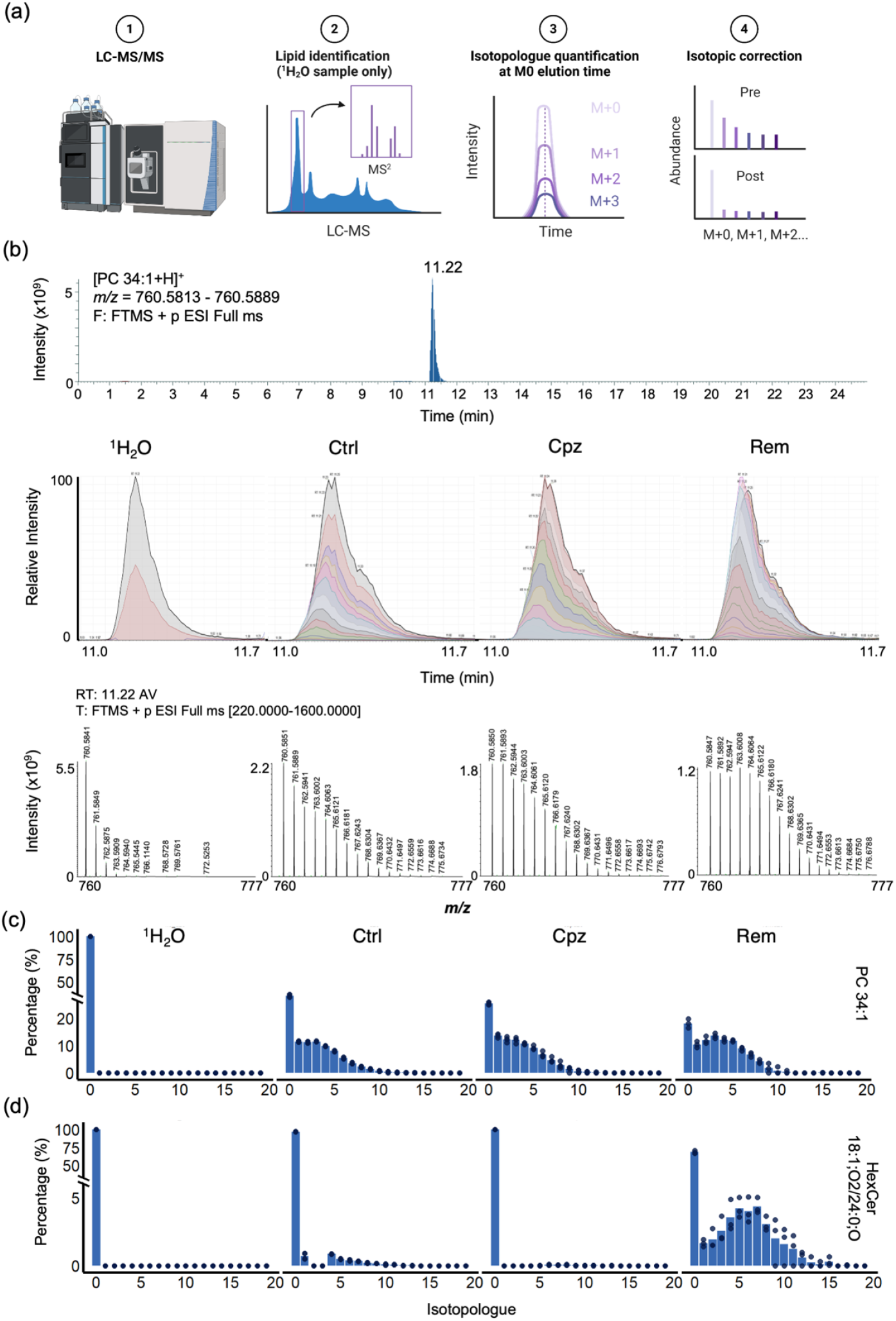
Quantification of deuterated isotopologues. a) Summary workflow for quantification of deuterated isotopologues. Following LC-MS/MS analysis of all samples, a list of the most abundant lipids in corpus callosum samples from mice administered only drinking water (^1^H_2_O control) was constructed based on MS/MS spectra and HPLC elution times. Isotopologues of each lipid were then quantified using the integrated peak area of chromatograms for lipid ions within 10 ppm mass tolerance of the theoretical *m/z* for deuterated isotopologues, and within a 0.1 min window of the elution time for the monoisotopic M+0 peak. Finally, peak areas were corrected for the abundance of naturally occurring heavy isotopes. b) Example mass spectra and chromatograms of non-deuterated and deuterated isotopologues of PC 34:1 from each experimental group. Upper panel: full-MS scan in positive ion mode at *m/z* 760.5851 ± 5 ppm from a ^1^H_2_O-only sample showing the PC 34:1 peak at 11.2 minutes. Middle panel: chromatograms of PC 34:1 isotopologues in a sample group from each treatment group, showing similar elution time window independent of deuterium incorporation. Bottom panel: full-MS^1^ spectra over the *m/z* range 760.0 – 777.0 at 11.2 minutes. c-d) Relative abundance of individual isotopologues from M+0 to M+20, expressed as a percentage of all isotopologues after isotope correction, for PC 34:1 (c) and HexCer 18:1;O2/24:0;O (d). Each data point represents one mouse.

### Identification of Lipid Biomarkers of Myelination

Applying these criteria, deuteration was quantified for 113 of the 171 identified lipids (Figure 3a). Robust deuteration above 25% was observed for 61 lipids in the corpus callosum of the chow-fed Ctrl mice, indicative of constitutive turnover in the two-week ^2^H_2_O administration period. These constitutively synthesised lipids were primarily comprised of phosphatidylcholine (PC, all twelve species), phosphatidylinositol (PI, all eight species), diacylglycerol (DG, five of six species), and triacylglycerol (TG, seven of eight species). In contrast, the myelin-enriched lipids including cholesterol, HexCer (all thirteen species), SHexCer (all six species), and most very-long chain sphingomyelins (SM, ten of thirteen species) showed minimal or no deuteration under basal conditions. This is consistent with the prior observation that turnover of lipids in myelin is slower for cholesterol and HexCer relative to PC and PE.^[26]^ Interestingly, seven of fourteen phosphatidylserine (PS) species were minimally deuterated under basal conditions. These minimally-deuterated PS species contained an 18:1 fatty acyl (FA) group paired with a very long-chain (≥20-carbon) saturated/monounsaturated fatty acid, a composition uniquely found in oligodendrocytes.^[32]^ Hence, the observed differences in baseline lipid deuteration likely reflect distinct cellular origins within the corpus callosum, with the more stable, long-lived lipids representing those produced by oligodendrocytes, and the rapid-turnover lipids representing those found in other neural cells, specifically neurons, astrocytes, and microglia.

**Figure 3.**
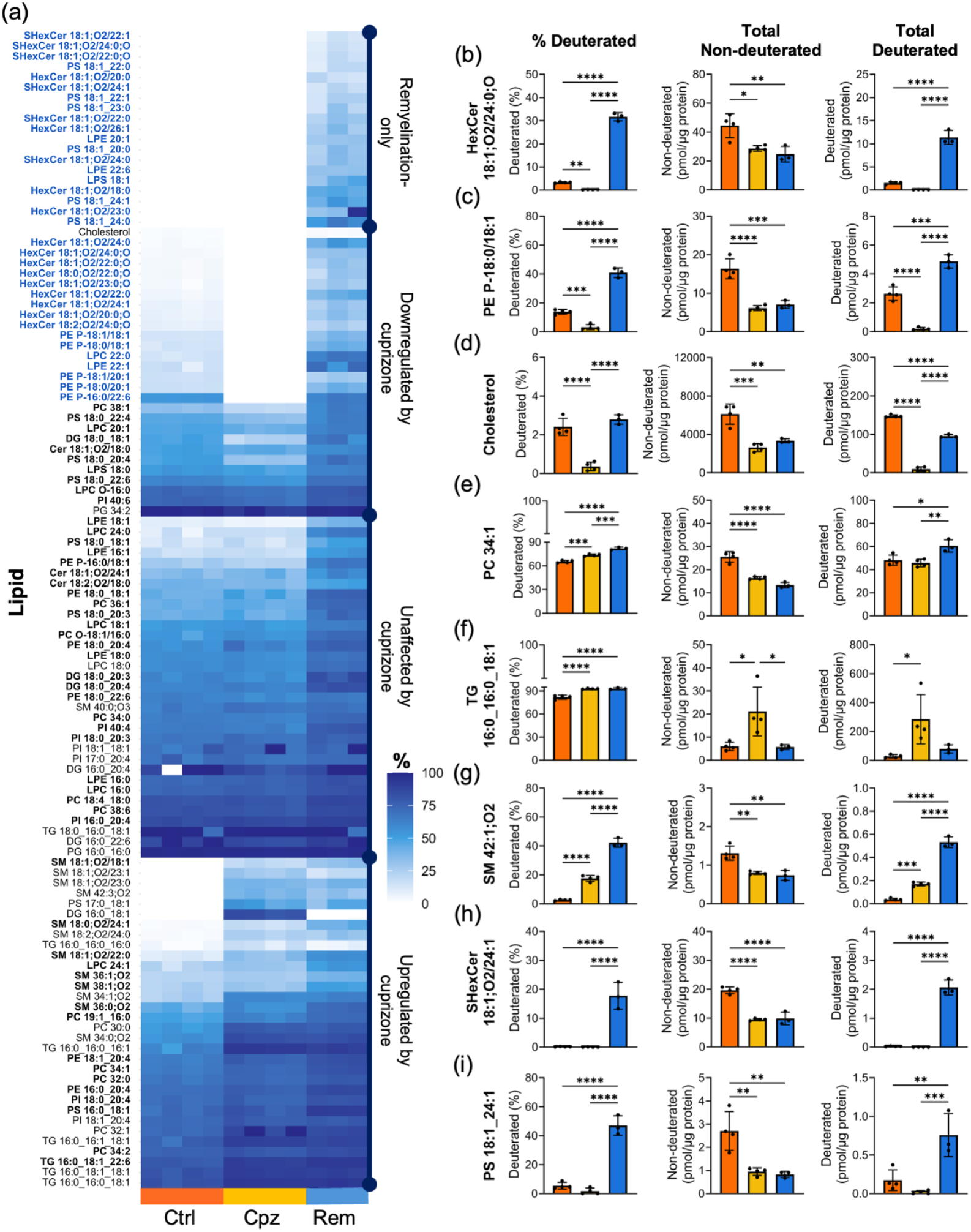
Identification of lipid biomarkers for myelination. a) Heat map of percentage deuteration for lipids defined as having quantifiable deuteration. Colour scale indicates percentage deuteration and each column represents one mouse. Data was not acquired for one of the four mice in the remyelination (Rem) group, due to a failed autosampler injection. Lipids with non-quantifiable deuteration in one or two of the experimental conditions were assigned a value of zero. Lipids were grouped into four categories based on changes in deuteration in response to cuprizone treatment and withdrawal. Lipid biomarkers for myelination (blue font) were identified based on the absence of deuteration during cuprizone intoxication, and a statistically-significant increase in deuteration relative to basal conditions during remyelination. (b-i) Percent deuteration, and total amounts of non-deuterated and deuterated HexCer 18:1;O2/24:0;O (b), PE P-18:0/18:1 (c), free cholesterol (d), PC 34:1 (e), TG 16:0_16:0_18:1 (f), SM 42:1;O2 (g), SHexCer 18:1;O2/24:1 (h), and PS 18:1_24:1 (i). Statistical significance was determined by one-way ANOVA with Tukey’s post-test for lipids with quantifiable deuteration in all three conditions, or by two-sample t-test for lipids with quantifiable deuteration in two out of three conditions. *p<0.05, **p<0.01, ***p<0.001, ****p<0.0001. Error bars represent standard deviation. p-values were corrected for false discovery rate, and lipids with significant increase in deuteration during remyelination relative to basal conditions at q<0.05 are in bold font in part (a).

The effect of cuprizone intoxication and withdrawal on lipid deuteration was assessed by two sample independent t-test for lipids with quantifiable deuteration in two conditions, and one-way ANOVA for lipids with quantifiable deuteration in all three conditions, adjusting p-values for false discovery rate. Cuprizone feeding significantly affected the deuterated proportion of 61 lipids (33 increased and 28 decreased). Consistent with the selective toxicity of cuprizone to mature oligodendrocytes and associated loss of myelin,^[33]^ the lipids showing decreased deuteration were mostly HexCer, PE-P, and cholesterol. The decrease in the deuterated proportion of these myelin-enriched lipids was accompanied by a decrease in the absolute abundance of their non-deuterated monoisotopes and almost complete absence of the deuterated form (Figure 3b-d). This reflects the loss of lipid that was present prior to ^2^H_2_O administration and lack of new lipid synthesis during cuprizone intoxication. Lipids that showed an increase in proportional deuteration during cuprizone feeding consisted of the more ubiquitous glycerophospholipids along with SM and TG (Figure 3a). Compared to basal conditions, these lipids were synthesised at a similar or higher rate during cuprizone intoxication (Figure 3e-g), consistent with prior findings for cuprizone- and lysolecithin-induced demyelination.^[34,35]^ Activation and proliferation of astrocytes and microglia are prominent features of the acute cuprizone model,^[36,37]^ and increased mRNA expression of Sphingomyelin Synthase 1 during demyelination has been observed in both cell types.^[35]^ Cuprizone intoxication also induces proliferation of oligodendrial precursor cells to replace the lost mature oligodendrocytes.^[30,36]^ Proliferation of these glial cells probably accounts for the observed increase in synthesis of particular lipids during cuprizone administration.

The withdrawal of cuprizone significantly altered the level of deuteration for 87 lipids and revealed a subset of myelination-specific lipids that were only deuterated in this treatment group. This group encompassed all the quantified HexCer and SHexCer, five of the six PE-P, six PS, three LPE, and one each of LPC and LPS. These myelination markers were characterised by an absence of deuteration during cuprizone-feeding, and robust induction of deuteration during remyelination to levels greater than baseline (Figure 3a). The increase in percentage deuteration was due to a large increase in the abundance of the deuterated lipid form (Figure 3b, c; h, i), indicating that *de novo* synthesis of these lipids supports myelin repair. Previous studies reported that levels of HexCer and SHexCer only show partial recovery during spontaneous remyelination, raising the concern that the lipid composition of remyelinated fibres does not recapitulate that of intact myelin.^[35,38]^ However, the pronounced acceleration in myelin lipid synthesis during remyelination observed herein suggests that a longer time frame may be needed for complete restoration of myelin, and determining rates of lipid synthesis over time will offer more detailed insights into remyelination dynamics.

Lysophospholipids are deacetylated forms of phospholipids generated from the enzymatic cleavage of ester bonds by phospholipase A1 and A2 enzymes.^[39,40]^ Thus the increase in deuterated lysophospholipids during remyelination likely stemmed from the hydrolysis of newly synthesised, deuterated glycerophospholipids. These species can be re-incorporated into membrane lipids through the Land’s cycle,^[40]^ or serve as lipid mediators to modulate inflammation to support local remyelination.^[34]^

Similar to other major myelin constituents, cholesterol synthesis was significantly increased during remyelination compared to demyelination (Figure 3d; total deuterated); however, the rate of synthesis remained higher under basal conditions, consistent with previous findings that cholesterol synthesis is rate limiting for myelination.^[41]^ During demyelination, cholesterol released from degenerating myelin is taken up by phagocytic microglia and stored as cholesterol esters, which can subsequently be reverse transported during remyelination^[42,43]^. However, whether myelin repair after acute demyelination is driven by cholesterol recycling or *de novo* synthesis remains a subject of debate.^[43-45]^ As cuprizone withdrawal did not restore the level of pre-existing cholesterol (Figure 3d, total non-deuterated), our results suggest that local synthesis of new cholesterol is a major contributor to myelin repair. The pre-existing cholesterol from damaged myelin could be lost from the central nervous system through a locally compromised blood brain barrier during early demyelination,^[46]^ or actively eliminated through enzymatic conversion into 24S-hydroxy-cholesterol.^[45]^

### Deuterium is Incorporated Primarily into the Lipid Acyl Chains

To determine if lipids were deuterated on the lipid headgroup, fatty acyl chains, or both, deuteration of product ions was assessed for five high abundance precursor ions that underwent fragmentation during data dependent acquisition. Fragmentation of the M+5 isotopologues of PE P-18:0/18:1 and PE P-16:0/18:1 in negative mode yielded product ions corresponding to the headgroup-derived phosphoethanolamine and dehydroglycerol phosphoethanolamine, 18:1 fatty acyl chain, and neutral loss of the 18:1 acyl chain (Figure 4a, b; Table S3, Supporting Information). The dehydroglycerol phosphethanolamine fragment (*m/z* 196.0380) resulting from cleavage of both acyl chains is diagnostic of PE-P, as it does not occur for the isomeric alkyl linked PE-O.^[47]^ The only isotopologue observed for the PE headgroup was M+1, whereas isotopologues up to M+5 could be found for fragments corresponding to the 18:1 acyl chain and neutral loss of the 18:1 acyl chain, implying that both acyl chains are deuterated. The *m/z* of the 18:1 acyl chain isotopologues differed as expected according to the number of deuterium atoms, despite a consistent mass offset relative to the expected *m/z*, suggesting that deuteration occurred primarily via metabolic incorporation of deuterium atoms into the fatty acyl chains. The relative paucity of deuteration on the PE headgroup may reflect the much greater number of hydrogens in the fatty acyl chains.

**Figure 4.**
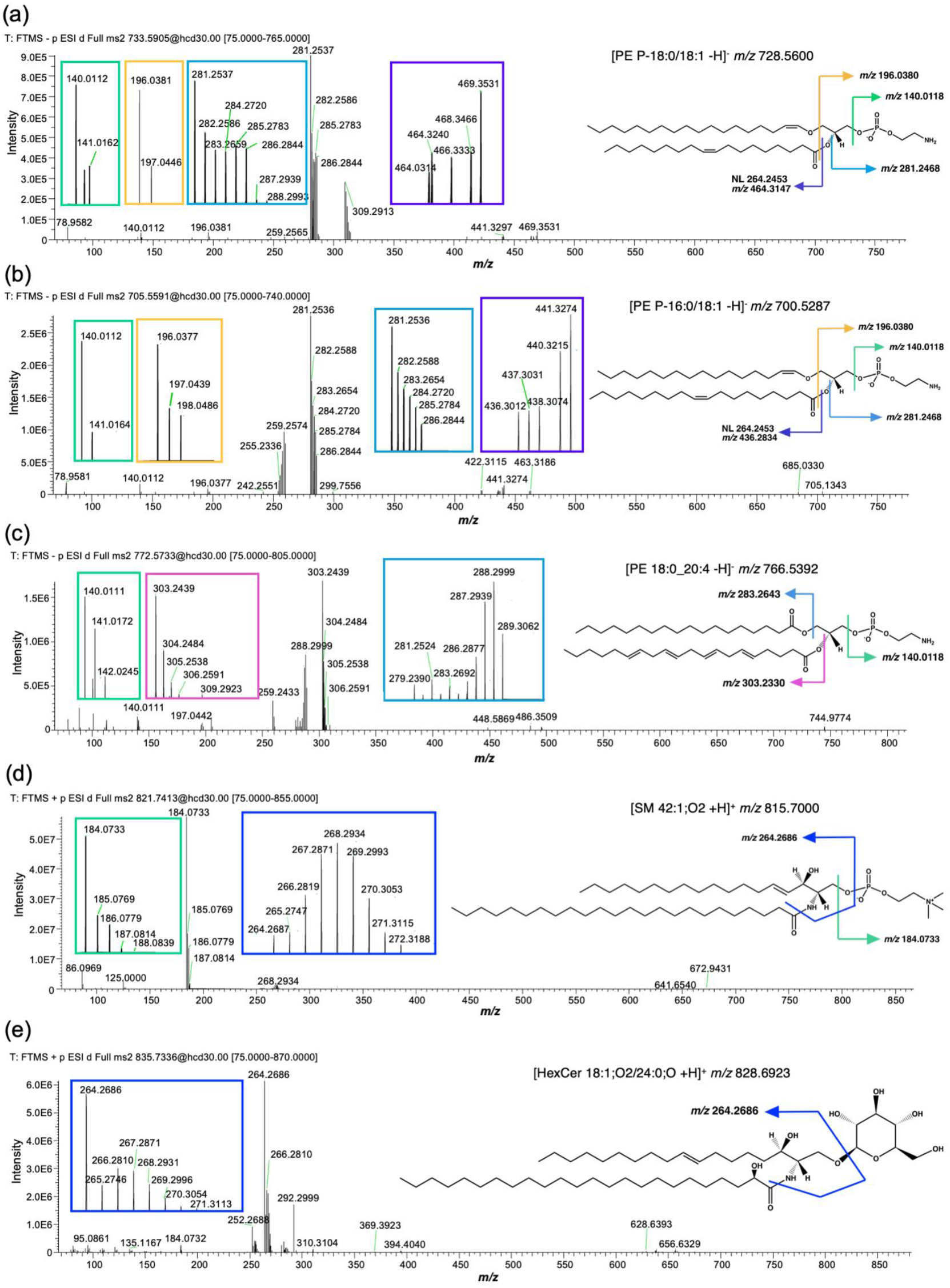
Deuteration occurs primarily on the lipid fatty acyl chains. Full MS^2^ spectra for M+5 PE P-18:0/18:1 (a), M+5 PE P-16:0/18:1 (b), M+6 PE 18:0_20:4 (c) in negative mode, and M+6 SM 42:1;O2 (d), M+7 HexCer 18:1;O2/24:0;O (e) in positive mode. Spectra were extracted from a sample from the remyelination group. Lipid structure and *m/z* of the non-deuterated precursor and fragments are shown on the right. Enlarged views of spectra corresponding to each fragment are shown in coloured boxes.

Fragmentation of the M+6 isotopologue of [PE 18:0_20:4 -H]^-^ yielded fragments with up to two ^2^H atoms for the PE headgroup, six for the 18:0 acyl chain, and three for the 20:4 acyl chain (Figure 4c; Table S3, Supporting Information). The most abundant isotopologues for the 18:0 acyl chain were M+4 and M+5, whereas M+0 was most abundant for the 20:4 chain, suggesting that the 18:0 chain was newly synthesised during the ^2^H_2_O administration period. In addition, deuteration appears variable for the same fragment in different lipids. Fragmentation of M+6 [SM 42:1;O2 +H]^+^ and M+7 [HexCer 18:1;O2/24:0;O +H]^+^ yielded the full range of possible isotopologues for the 18:1 sphingoid base, however the most abundant isotopologue of this fragment was M+4 for the SM, and M+0 for the HexCer (Figure 4d, e; Table S3, Supporting Information). Fragments corresponding to the choline phosphate headgroup in SM were indicative of both ^13^C and ^2^H atoms. The mass difference between the 185.0769 fragment and monoisotopic 184.0733 fragment is consistent with a ^13^C atom, while the adjacent peak at *m/z* 186.0779 is most consistent with one ^13^C and one ^2^H atom. As the resolution of the product ion scan used for this experiment does not resolve between the two isotopes, the detected *m/z* for these choline phosphate fragments most likely represent a mixture of [M+ ^13^C] and [M+ ^2^H] species.

### Synthesis of Myelin-Enriched Lipids is Localised to the Corpus Callosum During Remyelination

To map the localisation of newly synthesised lipids during remyelination, mass spectrometry imaging was applied to sagittal brain sections from the same mice as used for the above-described LC-MS/MS analysis. Tissue sections were pre-coated in non-endogenous lipid standards for signal normalisation and quantification before matrix application and analysis using atmospheric pressure matrix-assisted laser desorption and plasma post-ionisation.^[18]^ Due to the dense spectra lacking chromatographic separation of isobaric lipids and isotopologues, representative species [PC 34:1 +H]^+^, [PE P-34:1 +H]^+^, and [HexCer 40:1;O3 +H]^+^ were selected for analysis, as they exhibited minimal isobaric interference for their respective M+0 – M+6 peak clusters (Table S4, Supporting Information). Spectral peaks corresponding to the M+1 – M+3 isotopologues of [PC 34:1 +H]^+^ (Figure 5a, b), [PE P-34:1 +H]^+^ (Figure 6a, c), and [HexCer 40:1;O3 +H]^+^ (Figure 6b, d) were observed in all groups, including mice administered only ^1^H_2_O. Their presence in ^1^H_2_O control mice is most likely due to naturally-occurring ^13^C isotopes. Peaks corresponding to the M+4 – M+6 isotopologues of these lipids were only detected in mice administered 25% ^2^H_2_O.

**Figure 5.**
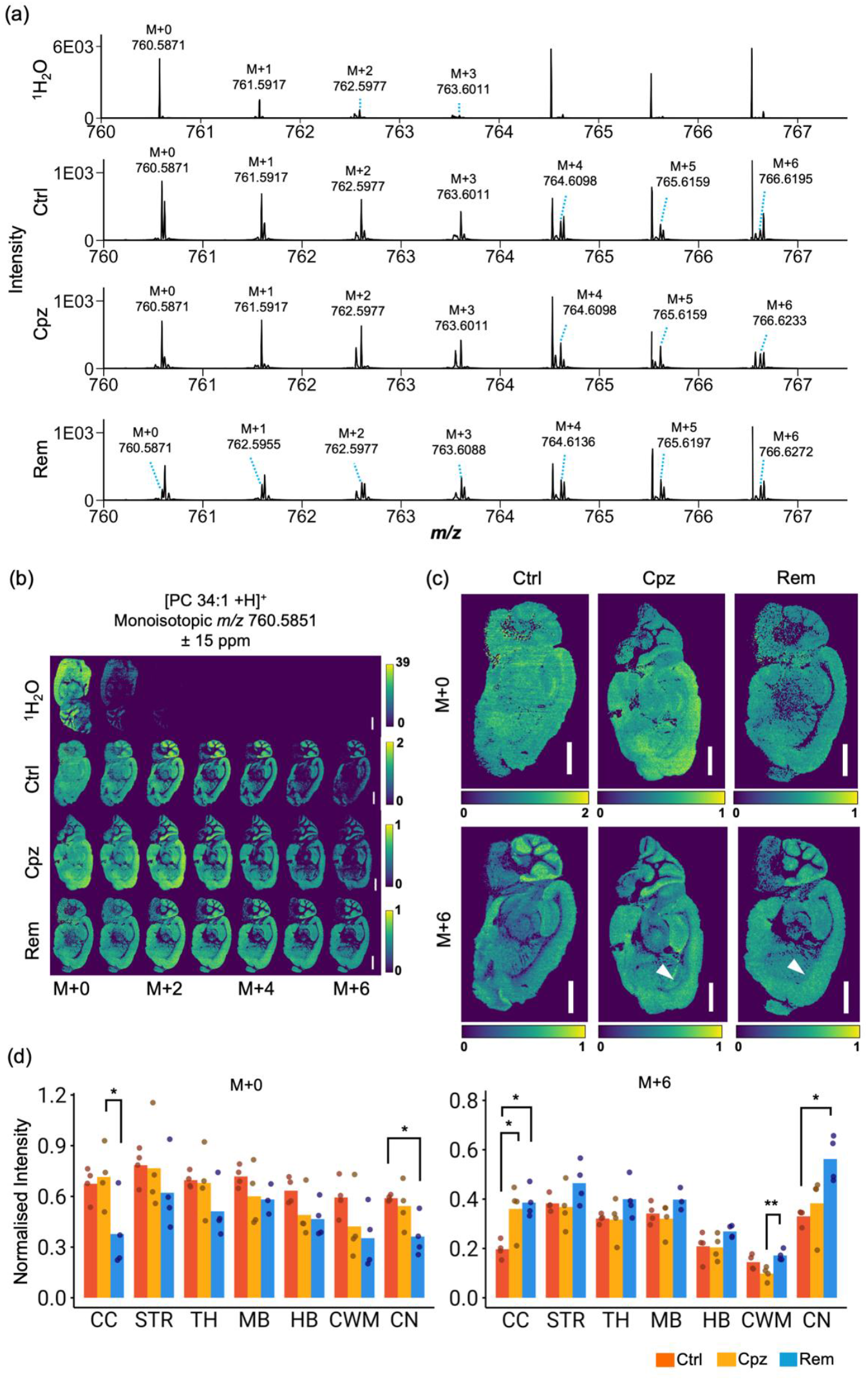
Visualisation of PC 34:1 isotopologues using MALDI-MSI. a) Full scan MALDI-MS spectra of the corpus callosum from *m/z* 760.0 – 767.0 showing the presence of deuterated [PC 34:1 +H]^+^. b) Corresponding ion images of [PC 34:1 +H]^+^ isotopologues from M+0 – M+6 extracted with a +/-15 ppm peak interval, and c) enlarged images of the M+0 and M+6 isotopologues. The colour scale represents the intensity of the ion of interest normalised to the [PC 33:1[D7] + H]^+^ internal standard on a per-pixel level. One representative image from each experimental group is presented. Scale bar: 2000 μm. Arrowheads denote increased M+6 localisation in the anterior corpus callosum during demyelination and remyelination. d) Mean normalised intensity of the M+0 and M+6 isotopologue in the corpus callosum (CC), striatum (STR), thalamus (TH), midbrain (MB), hindbrain (HB), cerebellar white matter (CWM), and cerebellar nuclei (CN). Statistical significance was determined by one-way ANOVA with Tukey’s post-test for each brain region: *p<0.05, **p<0.01. n = 4 mice per group.

**Figure 6.**
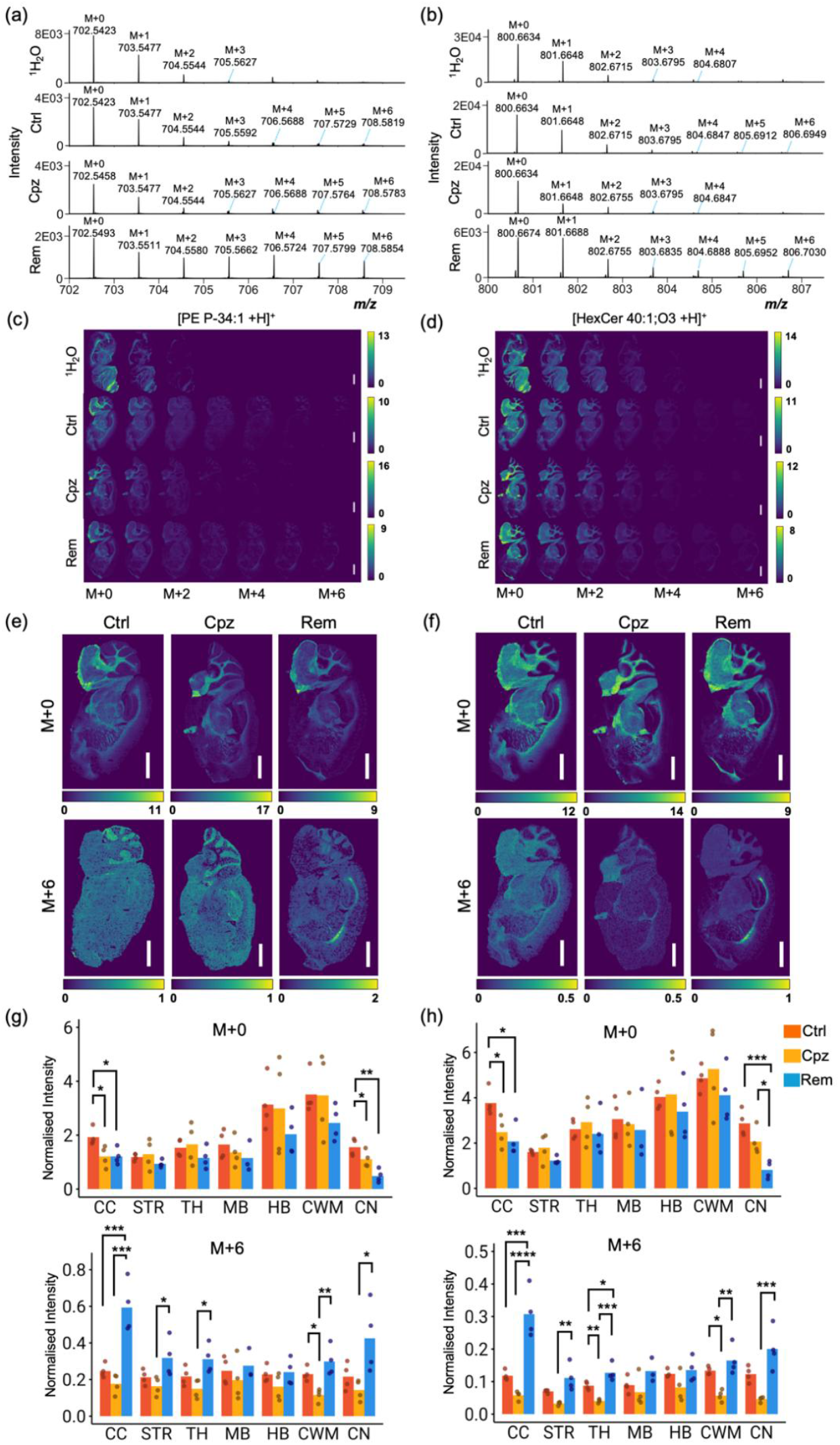
Newly-synthesised myelin lipids localise to the corpus callosum during remyelination. a-b) Full scan MALDI-MS spectra of the corpus callosum from *m/z* 702.0 – 709.0 (a) and *m/z* 800.0 – 807.0 (b) showing the presence of deuterated [PE P-34:1 +H]^+^ and [HexCer 40:1;O3 +H]^+^, respectively. c-d) Corresponding ion images of isotopologues from M+0 – M+6 extracted with a +/-15 ppm peak interval. e-f) Enlarged images of the M+0 and M+6 isotopologues. Colour scale represents the ion intensity as a ratio to the [PE 33:1[D7] +H]^+^ (e) or [HexCer 35:1;O2 +H]^+^ internal standard. One representative sample from each experimental group is presented. Scale bar: 2000 μm. g-h) Mean normalised intensity for the M+0 and M+6 isotopologues of PE P-34:1 (g) and HexCer 40:1;O3 (h) in the corpus callosum (CC), striatum (STR), thalamus (TH), midbrain (MB), hindbrain (HB), cerebellar white matter (CWM), and cerebellar nuclei (CN). Statistical significance was determined by one-way ANOVA with Tukey’s post-test for each brain region: *p<0.05, **p<0.01, ***p<0.001, ****p<0.0001. n = 4 mice per group.

In agreement with prior MSI studies,^[48,49]^ the M+0 isotopologue of PC 34:1 was distributed homogenously throughout the brain of Ctrl mice (Figure 5c). The M+2 – M+6 isotopologues of PC 34:1 were substantially more abundant in the grey matter regions, suggesting that basal turnover of PC 34:1 is slower in the white matter compared to the grey matter. In contrast, PE P-34:1 and HexCer 40:1;O3 were predominantly localised to the white matter tracts such as the corpus callosum, cerebellar and striatal white matter, and subcortical structures including the midbrain, hindbrain, and thalamus (Figure 6e, f). Consistent with the slow turnover of myelin, signals for the M+4 – M+6 isotopologues of these lipids were substantially lower than those of the monoisotopic species (Figure 6a-d).

To assess changes in pre-existing and newly-synthesised lipids during demyelination and remyelination, the abundance of the M+0 and M+6 lipid isotopologues were quantified in each of the annotated brain regions. Cuprizone treatment did not affect the M+0 or M+6 isotopologues of PC 34:1, apart from a localised increase in M+6 within the corpus callosum (Figure 5c, d). This observation aligns with our LC-MS/MS data indicating continued synthesis of PC 34:1 in this region during demyelination (Figure 3e), likely driven by proliferating glial cells. Following cuprizone withdrawal, the monoisotopic signal for PC 34:1 declined in the corpus callosum and cerebellar nuclei, consistent with the turnover of pre-existing PC 34:1 and its replacement by newly-synthesised, deuterated lipid, as evident in increased levels of the M+6 form (Figure 5d).

Cuprizone feeding significantly reduced the monoisotopic signals of PE P-34:1 and HexCer 40:1;O3 in the corpus callosum, and PE P-34:1 in the cerebellar nuclei (Figure 6g, h). This is consistent with histological evidence showing that these regions undergo the most pronounced demyelination in this model (Figure 1b),^[29]^ and therefore exhibit the greatest demand for remyelination. A pronounced enrichment of the M+6 form of these lipids was observed in the corpus callosum during remyelination (Figure 6e-h), however levels of the M+6 form were also significantly increased in other myelinated regions, including the cerebellar nuclei, cerebellar white matter, striatum, and thalamus.

These results demonstrate the capability of combining ^2^H_2_O labelling with MSI to visualise sites of dynamic lipid synthesis, and by extension, new myelin synthesis *in vivo*. While remyelination was not apparent in the anterior corpus callosum by histological staining (Figure 1b), MALDI-MSI showed synthesis of new myelin lipids in this region, indicating that remodelling is occurring at the molecular level.

## Conclusion

This study demonstrates the use of ^2^H_2_O administration in combination with high resolution mass spectrometry to quantify and visualise new lipid synthesis in the mammalian brain. Our approach builds on previous spatial metabolomics studies using stable isotope tracers for central carbon metabolites,^[50]^ instead using ^2^H_2_O to enable unbiased mapping of lipid metabolism events in living organisms. We applied these methods to demonstrate the differential synthesis rates of brain lipid synthesis and specific locations of myelin lipid synthesis following a demyelinating insult. This is important for the discovery and evaluation of remyelinating therapeutics, which are needed to enhance functional recovery and clinical outcomes in people with demyelinating diseases. More broadly, the methods described herein can be applied to many biological settings that require quantification and localisation of small molecule synthesis over a defined timeframe in living organisms.

## Supporting Information

The authors have cited additional references within the Supporting Information.^[51-54]^

### Experimental Section

#### Mice

8-week-old male C57BL/6J mice were obtained from Australian BioResources (ABR, Mossvale, NSW) and kept under a 12 h light/dark cycle with *ad libitum* access to food and water. Mice were monitored at least twice weekly. Experiments followed the Australian Code of Practice for the Care and Use of Animals for Scientific Purposes and were approved by the University of Sydney animal ethics committee (2022/2133).

At 10 weeks of age, mice were randomly assigned to receive standard chow pellets (Ctrl, n=4), pellets containing 0.2% (w/w) cuprizone (Merck #C9012) for 5 weeks (Cpz, n=4), or cuprizone pellets for 5 weeks, followed by standard chow for 2 weeks (Rem, n=4). ^2^H_2_O was purchased from Merck (#151882), filter-sterilised, and provided at 25% (v/v) in the drinking water during the final 2 weeks of the diet. A fourth group received drinking water only while on chow (n=1), cuprizone (n=1), or chow following cuprizone (n=2).

#### Brain Tissue Preparation

Mice were anaesthetised via isofluorane inhalation, then transcardially perfused with 0.9% saline. Brains were collected and halved sagittally. Corpus callosum tissue was dissected from one hemisphere using a 1 mm dissecting matrix and stored at -80°C. The other hemisphere was snap-frozen on dry ice. Sagittal sections of 15 μm thickness were prepared from the frozen hemisphere using a Shandon Cryotome FSE cryostat (Thermo Fisher Scientific), mounted onto Superfrost plus microscope slides (Thermo Fisher Scientific), and stored at - 80°C.

#### Lipid Extraction

The dissected corpus callosum tissue was homogenised in 500 μL of 20 mM N2-hydroxyethylpiperazine-N-2-ethane sulfonic acid (HEPES) buffer, pH 7.4, containing 10 mM KCl, 2 mM Na_3_VO_4_, 5 mM NaF, and complete protease inhibitor cocktail (Merck #11836170001), using a QSonica ultrasonication system at 4 °C, with 70% amplitude, 30 s on/off cycle for 10 min. The protein concentration of the homogenates was measured by bicinchoninic acid assay (Thermo Scientific #23225).

Lipids were extracted from 100 μL of corpus callosum homogenate using a two-phase methyl-tert-butyl-ether (MTBE)/methanol/water protocol.^[51]^ Homogenates were mixed with 850 μL MTBE and 250 μL methanol containing internal standards: 30 nmoles β-Sitosterol; 5 nmoles each of PC 19:0/19:0, TG 17:0/17:0/17:0, and CE 17:0; 2 nmoles of SM 18:1;O2/17:0, GlcCer 18:1;O2/17:0, PS 17:0/17:0, PE 17:0/17:0, and PG 17:0/17:0; PI 15:0/18:1[D7] and DG 15:0/18:1[D7]; 0.5 nmoles of SHexCer 18:1;O2/17:0, Cer 18:1;O2/17:0, LacCer 18:1;O2/12:0, LPC 17:0, LPE 17:1, and LPS 17:1 (Table S5, Supporting Information). Samples were sonicated for 30 min in a 4 °C water bath. Phase separation was induced by the addition of 112 μL milliQ water. Samples were vortexed and centrifuged at 2000xg for 5 min, and the upper organic phase was collected in 5 mL glass tubes. The remaining aqueous phase underwent two additional extractions using 500 μL MTBE and 150 μL methanol followed by sonication for 15 min and phase separation with 125 μL water. Organic phases from the three extractions were combined and vacuum-dried overnight in a Savant SC210 SpeedVac (Thermo Fisher Scientific). Dried lipids were reconstituted in 200 μL of 25% HPLC grade methanol/25% 1-butanol/50% milliQ water containing 0.1% formic acid and 10 mM ammonium formate, centrifuged at 2000 x g for 10 min to pellet the insoluble material, and 150 μL was transferred to glass HPLC vials.

#### Liquid Chromatography-Tandem Mass Spectrometry

Lipidomic data was acquired using a Thermo Fisher Q-Exactive HF-X mass spectrometer coupled to a Vanquish HPLC.^[51]^ Lipids were resolved on a Waters C18 Acquity UPLC column (2.1 × 100 mm, 1.7 μm pore size). The mobile phase flow rate was 0.28 mL/min using a 25 min binary gradient: 0 min, 80:20 A/B; 3 min, 80:20 A/B; 5.5 min, 55:45 A/B; 8 min, 35:65 A/B; 13 min, 15:85 A/B; 14 min, 0:100 A/B; 20 min, 0:100 A/B; 20.2 min: 80/20 A/B; 25 min: 80:20 A/B. Mobile phase A consisted of 10 mM ammonium formate, 0.1% formic acid in acetonitrile:water (60:40); mobile phase B consisted of 10 mM ammonium formate, 0.1% formic acid in isopropanol:acetonitrile (90:10). Data was acquired in full scan/data-dependent MS^2^ mode (resolution 60,000 FWHM, scan range 220–1600 *m/z*, AGC target 3e^6^, maximum integration time 50 ms) in both positive and negative mode for each sample. The ten most abundant ions in each cycle were subjected to MS^2^ using resolution 15,000 FWHM, isolation window 1.1 *m/z*, AGC target 1e^5^, collision energy 30 eV, maximum integration time 35 ms, dynamic exclusion window 8 s. An exclusion list of background ions was based on a solvent blank. An inclusion list of the [M + H]^+^, [M + NH_4_]^+^, and [M - H]^−^ ions for all internal standards was used. Positive mode data was not acquired for one sample from the remyelination group, due to a failed injection.

#### Identification and Quantification of Deuterated Lipids

Lipids from mice given normal drinking water (100% ^1^H_2_O) were annotated using LipidSearch v4.2 (Thermo Fisher Scientific), based on accurate precursor (5 ppm mass tolerance) and diagnostic product ions (8 ppm). A list of lipids, with their monoisotopic *m/z* and retention time, was generated (Table S1, Supporting Information). The predicted precursor *m/z* of the deuterated isotopologues was calculated by adding 1.0062767459 for each ^2^H atom (up to 20). Peaks for the non-deuterated monoisotope (M+0) and deuterated isotopologues (M+X, where X = number of deuterium atoms) in each sample were integrated using TraceFinder V5.1 (Thermo Fisher Scientific) based on precursor *m/z* (10 ppm window) and elution time (±0.1 min of the peak from the ^1^H_2_O control mice). IsoCor V2 (Python) was used to correct peak areas for naturally occurring heavy isotopes. The relative abundance of each isotopologue was calculated by dividing its peak area by the sum of all isotopologue peak areas (including the M+0 monoisotope) for that lipid. Deuteration was defined as having at least four adjacent isotopologues, with each isotopologue detected in at least three out of the four samples for any of the three treatment groups. Lipids were excluded from further analysis if the relative abundance of the M0 monoisotope was <95% in ^1^H_2_O control mice (i.e. if apparent deuterated isotopologues comprised ≥5% of total peak area for that lipid in ^1^H_2_O control mice, indicative of interference).

Percentage deuteration was defined as the summed peak area of all deuterated isotopologues (M+1 – M+20) expressed as a percentage of the total peak area for that lipid (sum of M+0 – M+20). Absolute amounts of each isotopologue in each sample were estimated by dividing the peak area by that of the class-specific internal standard, and multiplying by the amount of internal standard added. Lipid levels were expressed relative to protein concentration of the homogenate (pmoles lipid/μg protein).

#### Mass Spectrometry Imaging (MSI)

Sagittal sections between levels 10-11 were used for MSI. Following removal from -80°C, tissue sections were dried for 15 min in a vacuum desiccator. MSI SPLASH mix (Avanti Polar Lipids, Birmingham, AL, USA, #330841) was prepared by diluting the stock solution 10 times with LC-MS grade MeOH and sprayed onto the sections using a HTX-TM Sprayer (HTX Technologies, Chapel Hill, NC, USA) as detailed previously,^[52]^ before finally coating in 20 mg of 2,5-dihydroxyacetophenone (Sigma-Aldrich, Castle Hill, NSW, Australia) matrix using an in-house built sublimation system. Sublimation was performed at 140 °C for 2.5 min. Samples were then recrystalised at 50°C using 1 mL of 0.5% ethanol for 90 s and stored in a vacuum desiccator until MSI.

MSI was performed using a prototype timsTOF Pro mass spectrometer (Bruker Daltonics, Bremen, Germany). To enable MSI, the system includes an atmospheric pressure matrix-assisted laser/desorption ionisation (MALDI) ion source that is combined with an inline dielectric barrier discharge plasma ionisation system (SICRIT, Plasmion GmbH, Augsburg, Germany) as detailed previously.^[18]^ MSI was performed in positive ion mode at a pixel size of 30 × 30 μm^2^ using beam scanning dimensions of 15 × 15 μm^2^. The laser was manually focused onto the sample prior to MSI and operated at a repetition rate of 5 kHz with 200 laser shots accumulated at each position. Desorbed species were collected in the heated inlet capillary (middle temperature of 360°C) and transferred to the SICRIT device for plasma post-ionisation which was operated at 1,500 V amplitude and 15,000 Hz frequency.

All data were processed using SCiLS Lab 2026a (14.00.17781, Bruker Daltonics, Bremen, Germany). The luxol fast blue images were manually annotated in QuPath 0.6.0 to define regions of interest,^[53]^ which were imported using the SCiLS Lab annotation plugin for QuPath. The average peak area at selected *m/z* values (± 15 ppm interval width, centroided mass errors typically less than 5 ppm) were normalized to their respective class-specific internal standard and annotated regions were exported as a CSV file for visualisation and statistical analysis using R Studio version 4.2. Given the limited resolving power of the orthogonal TOF detector, M+2 deuterated lipid ions are often not separable from lipid ions with one less double bond (i.e. type II isobaric interference), thus the discussed lipids were selected based on inspection for minimal isobaric interference and spatial correlation between isotopologues. Ion images were produced by extracting the raw ion images for selected masses and normalising per pixel to the respective internal standard using the SCiLS API, Python 3.13 and NumPy 2.0.2, and were converted into CSV files for visualisation using FIJI.^[54]^

#### Luxol Fast Blue Staining

Tissue sections were stained with luxol fast blue (LFB) and cresyl violet after MSI. The slides were washed in 100% methanol for 30 seconds to remove matrix, after which the tissue was rehydrated using 95% then 70% aqueous ethanol, followed by Milli-Q water for 2 min each. Sections were fixed in 4% formaldehyde in PBS (Sigma-Aldrich, #HT501128) for 10 min and left to air-dry for 30 min inside a fume hood. Slides were placed in 0.3% HCl (prepared from 37% HCl, Sigma-Aldrich, #258148) in absolute ethanol for 3 min, rinsed for 3 min in 95% ethanol, and immersed in 1 mg/mL Luxol fast blue in ethanol (Sigma-Aldrich, #S3382) for 16 h at 58°C. Sections were allowed to equilibrate to room temperature and excess LFB washed with 95% ethanol, followed by a brief rinse in Milli-Q water. Slides were then immersed in a 0.05% lithium carbonate solution (Sigma-Aldrich, #255823) for 20 seconds to differentiate the sections followed by rapid, successive washes in 70% ethanol and Milli-Q water. This was performed for up to 1 min, until white matter areas were well-defined in blue against a clear grey matter background, and slides placed in Milli-Q water. The sections were counterstained with 1 mg/mL cresyl violet acetate solution (Sigma-Aldrich, #C5042) at 50°C for 20 min, then given a quick rinse in distilled water and differentiated in 95% ethanol for 3 min. Slides then underwent two changes of 100% ethanol for 2 min each and were cleared in xylene. Slides were cover slipped using DPX mountant (Sigma-Aldrich, #06522), dried overnight, and imaged with an Olympus Slideview VS200 in brightfield mode at 40x magnification.

#### Statistical Analysis

Statistical analyses were conducted using R Studio version 4.2. Lipids showing deuteration in two out of the three experimental groups were analysed by two sample independent t-test. Lipids showing deuteration in all three experimental groups were analysed by one-way ANOVA, followed by Tukey’s multiple comparisons test. p-values from both tests were binned and corrected for false discovery rate using the Benjamini-Hochberg test.

## Data Availability

The LC-MS/MS dataset supporting the conclusions of this manuscript is available in Tables S1, S2, and will be made available through Metabolomics Workbench. The MSI raw data files will be made available through Zenodo (DOI: 10.5281/zenodo.17504677).

## Acknowledgements

This research was supported by Ideas grants 2002660 (ASD) and 2028164 (ASD, SRE) from the National Health and Medical Research Council of Australia, Multiple Sclerosis Australia project grant 20-0113 (ASD), a Research Training Program scholarship (CZ), and Australian Research Council Future Fellowship FT190100082 (SRE).

The authors gratefully acknowledge use of the Imaging Facility, University of Wollongong; and subsidised use of the Sydney Mass Spectrometry and Laboratory Animal Services core facilities at The University of Sydney.

## Table of Contents

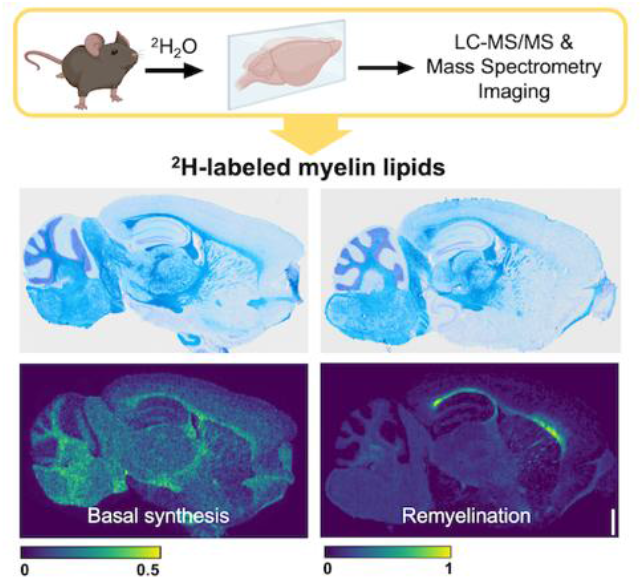

Liquid chromatography tandem mass spectrometry and mass spectrometry imaging were used in combination with deuterium oxide administration to quantify and localise newly-synthesised myelin lipids in the mouse brain. This methodology, used here to show sites of myelin repair, enables the measurement of dynamic lipid synthesis in living systems and spatial mapping of specific lipid metabolism events in complex tissue environments.

